# Higher-Order Biomarkers Through Network Motif Mining: A COVID-19 Case Study

**DOI:** 10.1101/2025.02.27.638877

**Authors:** Nicholas Chahley, Armstrong Murira, Nardin Nakhla, Carlos Oliver

## Abstract

We introduce a novel approach for analyzing expression data by integrating patient-level expression profiles with a Protein-Protein interaction network from the STRING database. Our pipeline leverages motif mining to identify recurring sets (motifs) of interacting biomolecules characterized by specific expression patterns, providing deeper insights into underlying biological processes. We applied our method to a publicly available dataset of plasma protein measurements from patients with mild/moderate COVID-19 and compared the motif features to those found by conventional differential expression analysis. Motif features demonstrated better performance in classification models and hierarchical clustering. Of note, they were able to resolve interpatient variability during clustering, while traditional features failed to do so. Interestingly, these discriminatory performances were achieved using a smaller and largely different set of proteins. Motif mining is a highly flexible method with capacity to integrate multiple modes of data and presents an exciting line of analysis for biomarker discovery as well as general biology.

## 1 Introduction

Since declaration of COVID-19 as a global health emergency in 2020, there have been over 7 million fatalities to date with a staggering 774 million confirmed cases [1]. While there have been significant gains made through robust vaccine efforts as well as therapeutics such as monoclonal antibodies and antivirals, an unresolved challenge is the varied trajectory of outcomes experienced upon infection. These range from out-patient hospitalization and fatal acute respiratory distress to short and long-term systemic impacts that are observed in patients with varying severity, even in the absence of well-defined pre-existing conditions (e.g., such as cancer, cardiovascular disease, diabetes, and obesity.) [2, 3, 4, 5] Furthermore, a lingering concern after infection is the progression to post-acute sequelae of SARS-CoV-2 infection (PASC), also termed ’Long COVID’; this condition is marked by persistent symptoms such as respiratory distress, cognitive dysfunction, and musculoskeletal pain, with a prevalence of 10% - 30% observed among the infected population[6]. In addition to pathogen-host interactions, many of these symptoms are likely attributable to aberrant and idiosyncratic immune-inflammatory responses that disturb pulmonary, cardiovascular, neurological, and hematological microenvironments. The incomplete characterization of the causal mechanisms behind these phenomena poses significant challenges in the realms of healthcare provision, public health policy, and pharmaceutical development [7]. Such obscurities in pathophysioloigcal phenomena are a primary motivation to discover biomarkers that accurately capture disease mechanisms.

It is recognized that most disease states arise from perturbations of a complex network of interacting biomolecules, rather than disregulation of a single gene or protein[8, 9]. These relationships can be represented as a network (or graph) where each biomolecule is a node and biomolecules which interact are connected by an edge. Additional metadata about biomolecules (e.g., subcellular location) and interactions (e.g., an inhibitory relationship) can be integrated as node and edge features, respectively, further enriching the data. Multiple mature, open-access, meta-resources allowing one to examine a biomolecule in its network context exist, for example: STRING[10] (protein-protein interaction), RNAInter[11] (RNA interactome), and HMDB[12] (metabolites interactome). Networks can then be overlaid with patient-specific data to create personalised models[13].

Yet the search for serum protein biomarkers remains widely focused on correlating the expression of one or more single molecules[14, 15], while failing to account for the context of protein interactions that more truly comprise a physiological response. Notably, regulation within and between cascades may occur through one or more intermediaries and interpatient differences in these regulatory elements could account for differing disease presentations or responses to treatment. Expression levels of a chain of interacting proteins provides a richer context of any pathology or therapeutic-intervention-driven changes. Here, treated as a single feature, such protein motifs could be used as higher-order, data-enriched biomarkers.

This paper details the in-depth examination of higher-order biomarkers in plasma protein profiles in COVID-19 patients who have exhibited mild to moderate symptoms. Using publicly-available next-generation plasma profiling data generated from 50 patients at the time of diagnosis and infection clearance [16] and the STRING protein-protein interaction (PPI) network, we have employed a novel approach that mines for subgraphs of interacting proteins and compares their frequency across samples to identify recurring patterns of expression within physically-interacting proteins. When combined with the highly multiplexed circulating proteome profiles, this approach allows for the identification of nuanced protein alterations that may correlate with disease progression and patient recovery. Here we show that our method, which takes into account combinations of proteins, more accurately discriminates patient cohorts based on their disease status in spite of interpatient variability among protein expression profiles. Furthermore, this approach reveals key proteomic signatures that distinguish disease and post-infection recovery states more accurately, highlighting potential underlying biological mechanisms at play during disease progression.

## 2 Methods

### 2.1 Data Source

Proteomics data were sourced from a previous study of COVID-19 patients with mild to moderate symptoms[16]. The dataset consists of 100 plasma protein samples taken from 50 patients each at two timepoints: first within 24h of having a positive COVID-19 PCR test (day 0, “disease phenotype”), followed by a second 14 days later and having a negative COVID-19 PCR test (day 14, “recovery phenotype”).

The protein profiles are measured as normalized protein expression (NPX) values, a log2 scale relative quantification unit. NPX data from 1459 protein variables (used as the input for motif mining) was obtained from Table S4[16]. The list of 239 differentially expressed proteins (used in the protein feature classification model) was taken from Table S6[16].

### 2.2 Motif Mining

We extracted protein signatures using Simmunome’s Omic Signatures™ pipeline. Omic Signatures™ extracts a list of protein-expression combinations (motifs) that are associated with a certain phenotypic characteristic. The pipeline uses a previously published method for motif mining (GASTON algorithm[17]). A motif is a set of three protein-level pairs that are directly linked in patients graphs (e.g., MAP2K5-low/RB1CC1-low/SIRT1-low and MAP2K5-high/RB1CC1- low/SIRT1-low are two distinct motifs). In the current study, GASTON was run with parameters support=30 and size=3.

To create the patients graphs, we organized the data into two groups (n = 50 each): disease and recovery. NPX values are binned into five expression levels (very low, low, medium, high, very high) based on a histogram of the pooled values of the entire dataset. A protein-protein interaction (PPI) graph for each patient is constructed using the STRING database[10] and their expression level data at each timepoint (Figure 1). More specifically, each patient graphs nodes correspond to a protein for which we have a NPX measurement and two nodes are connected when a physical interaction is annotated by STRING. The patients are randomly split into training and test subsets (80/20%) where both of a patient’s two graphs are exclusively in either set. Motif patterns are mined from the training set of each group, combined into a list, and their frequencies are counted across both groups. No motifs were generated from the test data.

**Figure 1:**
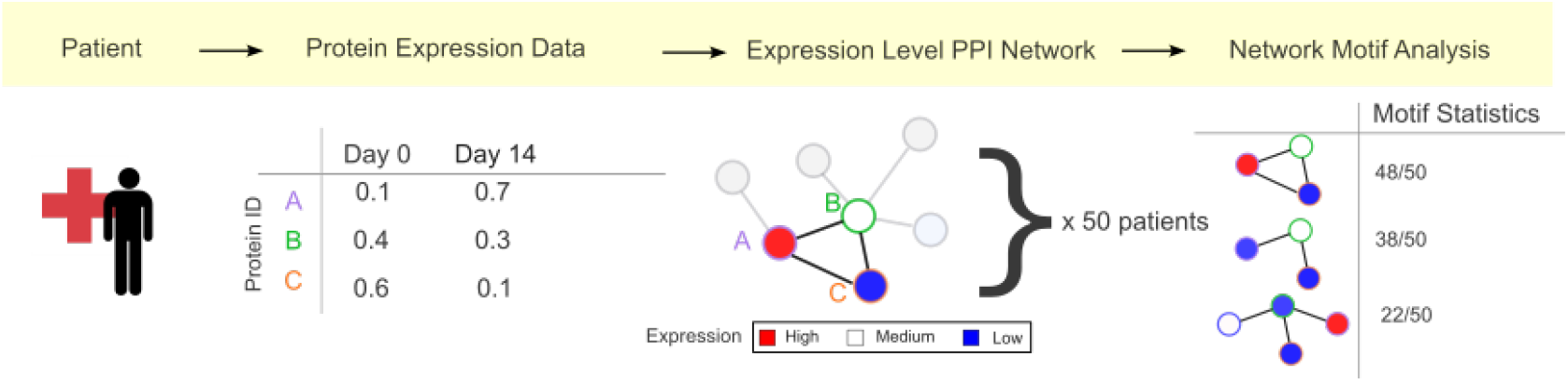
Motif mining schematic. A protein-protein interaction (PPI) network (based on STRING v12) is constructed from the expression level data for each patient at each timepoint (day 0 and day 14). Recurring motifs (sets of 3 protein-expression pairs) are identified within each timepoint.

Motifs enriched in each group are identified by frequency analysis. Motif frequency is counted in each group, and absolute difference between groups. Motifs are considered associated with the phenotype in which they have the highest frequency (e.g., a motif which occurs in 30 samples in the disease group and 10 samples in the recovery group is a disease-associated motif).

### 2.3 Statistics

#### Frequency statistics

We tested motifs for statistical significance by computing *χ*^2^ statistics on their absolute between-group differences. Family-wise error rate (FWER) was controlled to *p <* 0.05 using Bonferroni multiple testing correction.

#### Permutation ablation

To investigate the specificity of motifs to patient-groups, we apply permutation of patient group laels to estimate the frequency of the identified motifs in the population of pooled samples.

At each iteration, 40 samples are randomly selected from the total pool (n = 100), used as input for motif mining, and the counts of motifs mined from the real experimental groups (Section 2.2) with a group difference ≥ 0 are recorded. A two-tailed t-test is performed for each motif comparing the observed counts (in the disease and phenotype groups) and the synthetic mean (*X*_*phenotype*_ − *μ*_*synthetic*_)). FWER is controlled to *p <* 0.01 by Bonferroni correction.

Two experiments are conducted, repeating 1,000 and 10,000 times, respectively. All observed motif frequencies were higher and statistically significant (*p <* 0.01) compared to their synthetic means.

### 2.4 Information Encoding Experiments

To investigate whether higher-order motifs capture data variation more effectively than individual protein levels, we examined the effects of dimension reduction on both feature vectors. We compared the differentially expressed proteins from [16] (N=239) to the significant motifs we extracted from the same dataset (N=737). The motif profile feature for each sample is a vector of categorical variables indicating the presence or absence of a specific motif. Analyses are performed using the scikit-learn python library[18].

#### Dimensionality reduction

Numeric data (i.e. protein expression) is scaled to mean=0 and SD=1 and missing values are set to zero. Principal component rotation is fitted on the only training data to avoid data leaks. Principal component analysis (PCA) is performed in Python using sklearn.

#### Classification models

XGBoost models are generated on the training data using the protein and motif features in full, and after dimension reduction over a range of principle component numbers (PCNs). To evaluate the discriminatory capacity of their respective feature sets, models are given the task of predicting phenotype of the test data, given protein level measurements. Hyperparameters of each model are optimized for receiver operating characteristic (ROC) area under curve (AUC) score by a random search with two-fold cross validation targeting.

### 2.5 Phenotype Discrimination Experiments

#### Hierarchical clustering

Dissimilarity matrices are computed on Pearson’s distance for the protein features, as described in [16], and Jaccard distance for the motif features. Hierarchical clustering is performed using the Ward2 algorithm and dendrogram visualization using R packages ape and dendextend[19, 20].

## 3 Results

### 3.1 Motif Frequency and Composition

We identified 53 statistically significant (*FWER <* 0.05) motifs (from 2989 returned by subgraph mining) made up of 53 proteins (Table S1). These motifs ranged in frequency from 25–65 (median 33) and absolute group differences from 23–36 (median 27). Four proteins, LAG3, IFNG, GRN, and CD74, appeared at multiple levels. All present only at high level in disease-associated motifs and at medium level in recovery-associated motifs. The majority of proteins composing the statistically significant motifs are distinct from those identified through traditional differential expression analysis (Fig. 2).

**Figure 2:**
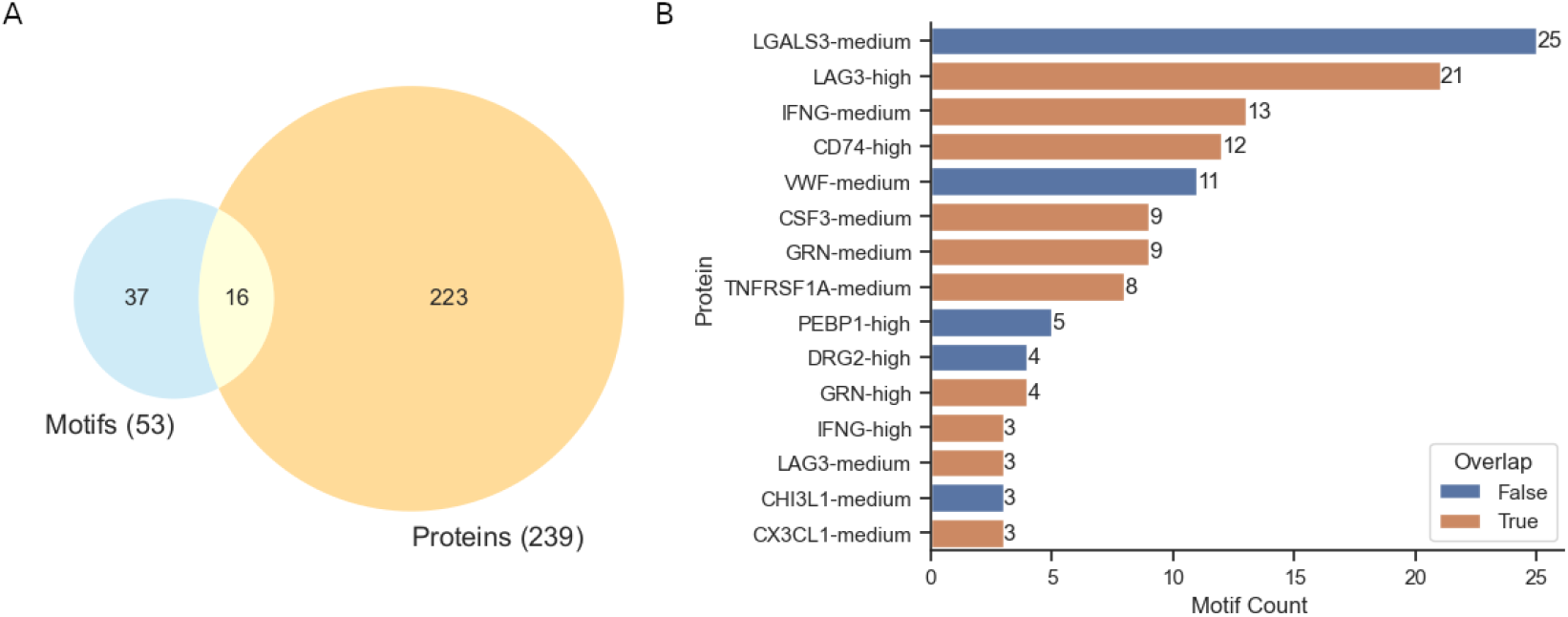
(A) Overlap of statistically significant proteins identified by traditional differential expression (“Proteins”) and motif mining (“Motifs”). (B) Most frequent motif proteins, ranked by number of statistically significant motifs they appear in. Overlap indicates a protein that is one of the 16 overlapping proteins identified by both methods. The full set of protein overlaps can be found in Table S2.

### 3.2 Information Encoding Experiments

#### Motifs better retain COVID-19 predictiveness at low Principle Component Numbers (PCN)

Unbiased cumulative variance is enriched in motif principal components, which reach a sum of 95% in fewer PCN than the protein features (7 vs 50, Fig. 3). Fig 4 compares the predictive performance of features reduced to the same dimension at various PCN (ranged around each feature’s 95% cumulative variance threshold) in a XGBoost decision tree classifier. Motifs produced classifiers with higher AUC scores than proteins at the same PCN over all 6 levels.

**Figure 3:**
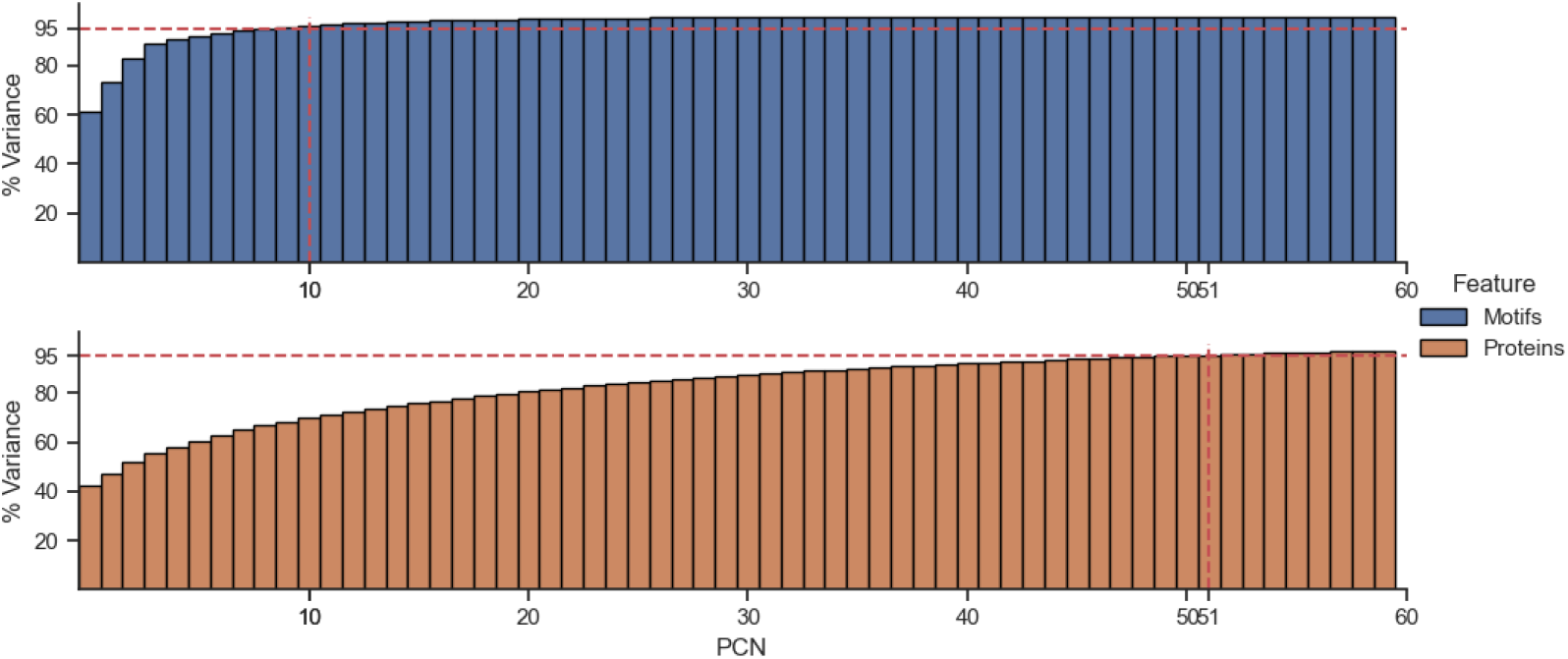
Cumulative variance for the first 60 principle components (PCN) of the significant motif (N = 53) and protein (N = 239) profiles. Dashed lines indicate the PCN which reaches 95% variance explained (10 and 51, respectively).

**Figure 4:**
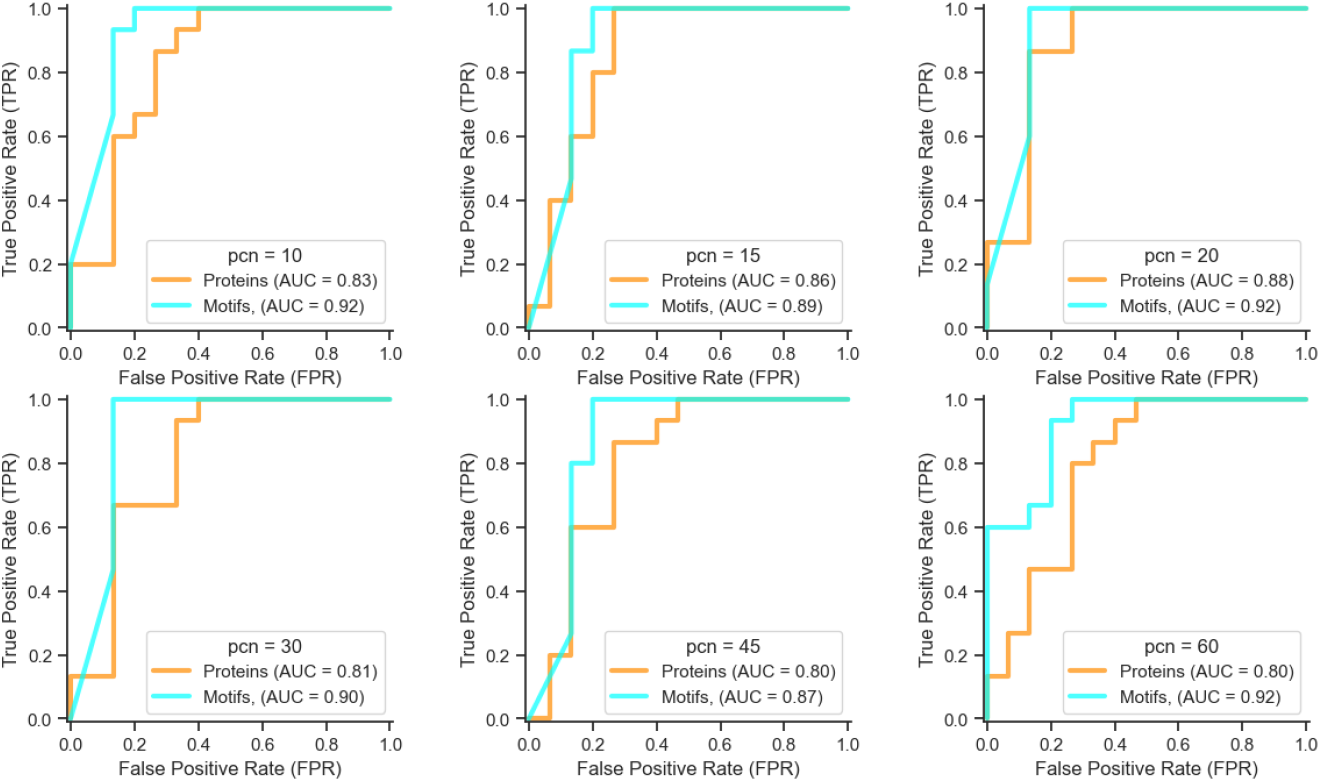
Receiver operating characteristic (ROC) curves obtained from the XGBoost models using protein (blue) or motif (red) features. Classification objective is binary logistic predicting phenotype (disease or recovery) probability on the test dataset (N=20).

### 3.3 Phenotype Discrimination Experiments

#### Motifs achieve sharper hierarchical clustering of COVID-19 status

Hierachical clustering results are visualised as dendrograms in Fig. 5; tip labels are colored by sample phenotype. Fig. 5A and Fig. 5B compare the motif approach of the current analysis with the protein approach originally taken[16]. We see a larger clusters of mostly disease (red; bottom-right vs middle-left) and recovery samples (blue; top-right vs bottom-left) from the motifs approach. This suggests a larger number of similar motif signatures per-group. The majority of participants’ protein signatures are most closely clustered with themselves (i.e. they have a mutual shortest cophenetic distance), compared to zero instances among motif signatures (Fig. 5C). This remains the case when the data is split into the same training and validation groups used for mining test (Fig. 5D). This suggests that there is less intersubject variability in motif profiles than protein profiles obtained from this dataset.

**Figure 5:**
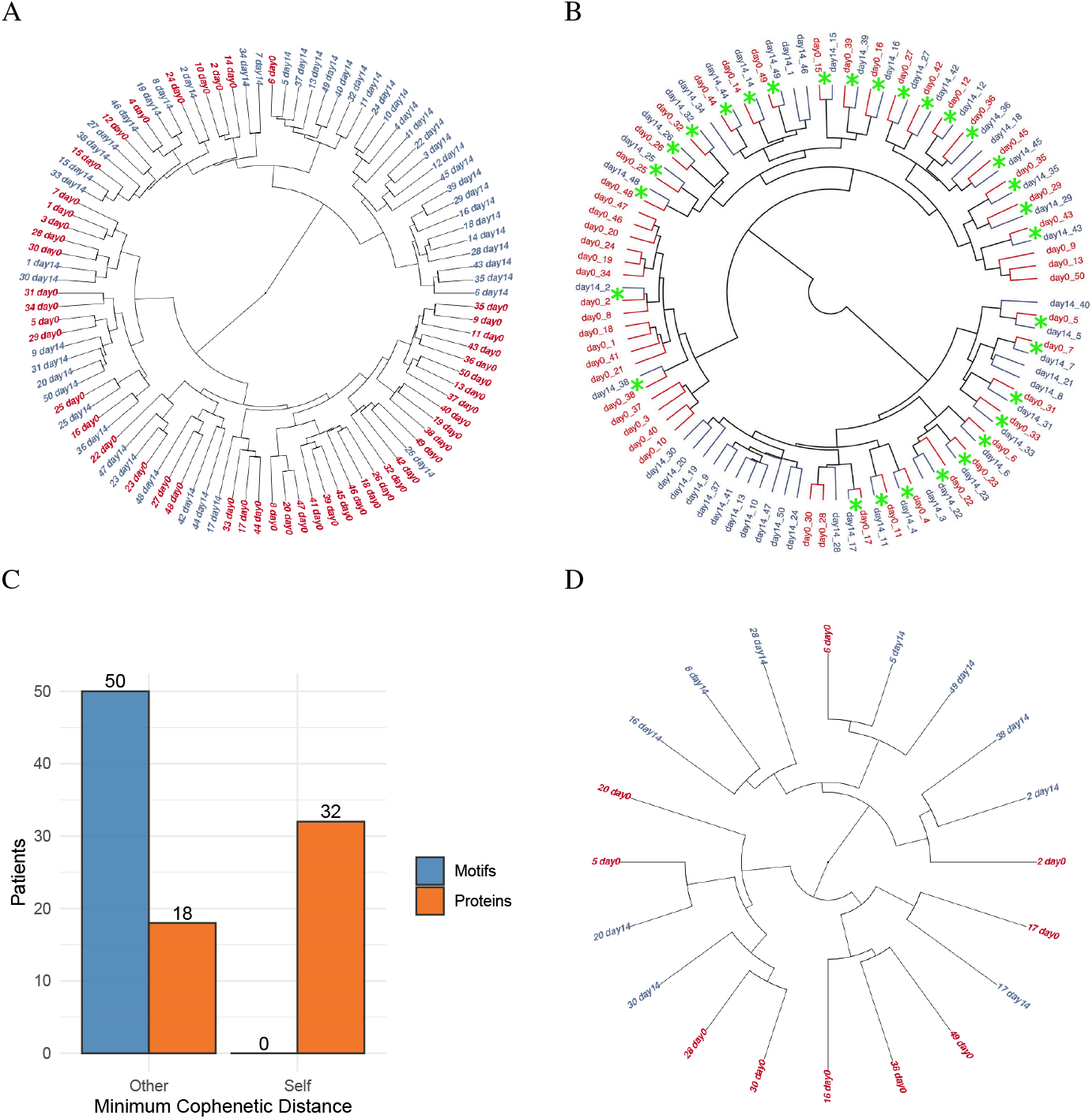
(A/B) Dendrograms from hierarchical clustering of all samples using motif presence (A) and differential protein expression (B – taken from [16]). (C) Count of instances where both samples from the same patient are most similar, i.e. they have a mutual shortest cophenetic distance (marked with green asterisks (*) on dendrograms). (D) Clustering of the test subset using motifs mined from the training subset. Samples are colored by phenotype (disease in red, recovery in blue).

## 4 Discussion

Discernment of biologically relevant biomarkers presents significant challenges, especially when considering the vast heterogeneity that exists between populations and individual patients. As such, discovery must account for this variability and ensure that biomarkers are robustly indicative of a particular disease state or therapeutic response across different genetic backgrounds and environmental exposures. This necessitates large-scale studies to differentiate between what might be a true, universal biomarker *versus* those that are only relevant in specific subpopulations or even singular patient cases. Importantly, in the case of limited biomarker presentation in a population or subpopulation, e.g., in the case of a rare disease, large scale studies are not feasible and as such, low-data methods without compromising robustness are necessary.

One of the major hurdles in this area is the enhancement of the signal-to-noise ratio. Omics technologies, which generate high-dimensional data, can often produce a significant amount of noise. Typically, dimensionality reduction techniques such as principal component analysis (PCA), t-distributed stochastic neighbor embedding (t-SNE), and others are critical in filtering out noise and focusing on the most informative features of the data. This is not only important for the clarity of the resulting data but also for the manageability and interpretability of the dataset.

However, even with clear signals, understanding the biology in context is paramount. The differential expression of a single protein or gene does not yield much information in isolation. Biological systems are complex and highly integrated, and thus, it is the interplay between various biological molecules and pathways that needs to be considered. This is where a network analysis approach becomes essential. By examining the neighborhood in which changes occur and quantifying the relative change in expression within two or more different conditions, researchers can gain insights into the pathways and processes that are truly impacted by or are indicative of a disease state. Network analysis helps to map out the interactions and understand the systemic effects of biological changes, rather than focusing on isolated elements which may not be as informative.

In our hands, we tackle both challenges of signal-to-noise ratio (SNR) as well as biological context for differential expression by employing a sophisticated technique of subgraph mining and frequency ranking as preliminary filtering steps. Here, we found that using protein motifs better captured the complexity in our dataset without losing significant information. Specifically, this approach explained 95% of the variance in the proteomic readout from the samples with fewer PCNs, demonstrating a more efficient and powerful method in comparison to the traditional unit protein approach (Fig. 3). Next, we assessed the accuracy of protein motifs vs unit proteins in predicting the disease vs. recovery phenotype on day0 and day14, respectively (Fig. 3). Here, dimensionality reduction for proteins was carried out using PCA whereas for motifs, this was conducted using multiple factor analysis. Similar to the variance analysis, we found that motifs were notably more accurate at predicting phenotype with fewer PCNs (Fig. 3b). Interestingly, with a higher number of PCNs in both proteins and motifs, we see a convergence in the AUC scores. Altogether, we surmise that by capturing 95% variance of the dataset and obtaining higher classifier AUC with fewer PCNs, motifs obtained from subgraph mining have a higher feature efficiency. In comparison to unit proteins, these motifs more efficiently capture the discriminatory features in phenotypically distinct biological/ clinical datasets, which is likely driven by the context of assessing natively interacting proteins. Conversely, with increasing PCNs, the convergence of AUCs indicates that as more components are included in the analysis, the additional components start to add less discriminative value. Here, both methods eventually capture enough of the dataset variance to provide a similar level of predictive accuracy. This can imply: (i) redundancy in the information captured by the additional PCNs beyond a certain point, whereby both methods are essentially capturing the same information or (ii) saturation whereby both approaches likely contain a core set of informative features, and beyond this core, additional features contribute less to improving model performance.

Furthermore, we demonstrated refined clustering of patient phenotypes when using the subgraphs, in contrast to the single protein approach (Fig. 5). In the latter, individual patient differences predominantly influenced clustering, indicating a less clear distinction between the two time points. Here the difference in performance is more stark compared to the classification task: while the single proteins achieve lower, but acceptable ROC curves with a trained classifier, they fail to overcome interpatient variability in unsupervised clustering. This supports the role played by subgraphs in aligning more closely to the different underlying biological processes in subjects at the two timepoints. Notably, our motifs currently utilize triples; however, the introduction of higher-order motifssuch as quadruples or quintuplesmay offer additional improvements to the SNR, potentially leading to even more distinct and reliable clustering outcomes.

It is interesting that only a small number of protein identities are shared between the significant features of the two methods (Fig. 2A). This suggests that, in addition to improved SNR, our method can provide value through highlighting novel features. There is evidence linking a number of the hub nodes uniquely identified by our method to COVID-19 (Fig. 2B, blue). For example: upregulation of LGALS3, which plays a role in the airway epithelial barrier, has been associated with COVID-19 positive bronchoalveolar lavage tests[21]. A meta analysis found VWF as a marker of COVID-19, as well as disease severity in conjunction with ADAMTS13 [22, 23]. CHI3L1 has been associated with mortality and clinical outcome and identified as a therapeutic target[24, 25]. Furthermore, the most frequent overlapping proteins also have established links to COVID-19, e.g., LAG3, IFNG, and CD74[26, 27, 28]

Future investigations should aim to dissect these subgraphs further, perhaps through targeted proteomics or functional assays, to validate their roles and interactions. Longitudinal studies that track the evolution of these subgraphs throughout the course of a disease and into the recovery phase would be particularly illuminating. Such detailed analyses could pave the way for subgraph-specific interventions, which may offer more precise therapeutic strategies than those targeting single genes or proteins. Furthermore, the dynamic interplay between various pathways during disease progression and subsequent recovery phases can offer potential biomarkers for disease prognosis and recovery. Moreover, they may provide a roadmap for the development of targeted therapies that modulate specific pathways at different stages of disease.

In summary, our study provides a refined perspective and a novel approach to better understand the complexity of the immune response in disease and recovery states through protein interaction subgraphs. This approach holds promise for uncovering the intricate molecular choreography of any biological systems in diverse conditions ranging from health, disease or under treatment. Overall this approach offers novel avenues for defining biomarkers or underlying biological mechanisms in the context of basic research through to therapeutic intervention in the realm of personalized medicine.

## Supporting information

Supplemental Table 1 - motifs

Supplemental Table 2 - node level counts

## Notes

### Competing Interest Statement

The authors have declared no competing interest.

